# Interactions between the neural correlates of dispositional internally directed thought and visual imagery

**DOI:** 10.1101/857490

**Authors:** Theodoros Karapanagiotidis, Elizabeth Jefferies, Jonathan Smallwood

## Abstract

Cognition is not always directed to the events in the here and now and we often self-generate thoughts and images in imagination. Important aspects of these self-generated experiences are associated with various dispositional traits. In this study, we explored whether these psychological associations relate to a common underlying neurocognitive mechanism. We acquired resting state functional magnetic resonance imaging data from a large cohort of participants and asked them to retrospectively report their experience during the scan. Participants also completed questionnaires reflecting a range of dispositional traits. We found thoughts emphasising visual imagery at rest were associated with dispositional tendency towards internally directed attention (self-consciousness and attentional problems) and linked to a stronger correlation between a posterior parietal network and a lateral fronto-temporal network. Furthermore, decoupling between the brainstem and a lateral visual network was associated with dispositional internally directed attention. Critically, these brain-cognition associations were related: the correlation between parietal-frontal regions and reports of visual imagery was stronger for individuals with increased connectivity between brainstem and visual cortex. Our results highlight neural mechanisms linked to the dispositional basis for patterns of self-generated thought, and suggest that accounting for dispositional traits is important when exploring the neural substrates of self-generated experience (and vice versa).

## Introduction

Cognition is not always focused on events in the here and now; we often engage in patterns of self-generated thoughts that are, at best, only loosely related to the events in the here and now. Contemporary accounts of self-generated thought have identified thematic patterns, or “dimensions” of thought [1], with a reproducible structure across studies and tasks contexts [2–4]. Recurring themes in experience capture variation in the content and form of our thoughts [5] and highlight stable patterns, such as thoughts about the future or the past [3, 6, 7], the modality of experience, i.e. whether it is dominated by imagery or inner speech [3, 8], or the level of subjective detail [3–5].

Using a variety of experience sampling techniques, such as the Experience Sampling Method [9], Ecological Momentary Assessment [10], and Descriptive Experience Sampling [11], as well as questionnaires, such as the Imaginal Processes Inventory [12], previous studies have shown that patterns of self-generated thought have associations with dispositional variables related to affect, attention, psychological well-being and disorders. For example, visual imagery and inner speech, two of the most often reported modalities of naturally occurring thoughts [13, 14], have both been related to multiple psychopathologies. Negative mental images can accelerate the onset of depression and lengthen the duration of an episode [15], while intrusive, recurrent, spontaneous imagery has been linked to psychiatric disorders, such post-traumatic stress disorder, obsessive-compulsive disorder, health anxiety and social phobia (see [16] for a review). Similarly, inner speech has been implicated in psychotic, mood, and anxiety disorders ([17] for a review). In addition to mental imagery, future thinking has been shown to improve mood [6] and facilitate problem solving by refining future goals [2]. In contrast, thinking about the past is associated with negative mood [6, 18] and in the case of intrusive, ruminative thoughts, to depression and anxiety [19]. Finally, identified patterns of spontaneous thought have been linked to creativity and intelligence [1, 20, 21] as well as ADHD [22], neuroticism [23] and measures of self-consciousness [24]. These studies demonstrate that different themes of ongoing thought may reflect dispositional variation along a range of dispositional variables.

Our study set out to understand how patterns of neural organisation at rest underpin the associations between dispositional traits and patterns of ongoing experience. It is often assumed that the default mode network (DMN, [25]) plays a particularly important role in self-generated thought (i.e. [26]). Consistent with this view, studies combining experience sampling with online neural measures have shown that core regions of the DMN, such as the posterior cingulate cortex (PCC) and the medial prefrontal cortex (mPFC), can have greater activity when patterns of ongoing thought shift from task relevant information ([27, 28], although see [4]). Studies suggest the PCC may be an integrative hub associated with the content and form of thoughts [3], while connectivity between the hippocampus and the mPFC has been linked to episodic contributions to ongoing experience [7]. Likewise, increased low frequency fluctuations in activity in the PCC have been associated with greater imagery [5]. However, studies have also found that patterns of default mode network connectivity are predictive of mindfulness [29], attention deficit disorder [30], depression [31], and life satisfaction and unhappiness [32, 33], each of which have associations with patterns of ongoing thought (for example, Mindfulness, [34]; ADHD, [22, 35]; Depression and unhappiness, [36]). Together, such evidence establishes that regions within the DMN are important in both patterns of ongoing experience, as well as many of the dispositional traits that are associated with aspects of ongoing experience within the general population.

More recent studies have highlighted neural processes outside the DMN that make an important contribution to ongoing cognition, either directly or indirectly by supporting dispositional traits that are linked to patterns of experience. Executive [28, 37] and attention systems [8, 38] have been related to variations in whether attention is focused on an external task. At the same time, neural patterns within the fronto-parietal system are linked to intelligence [39], a trait linked to the ability to maintain focus during complex tasks [40] and to limiting self-generated thoughts to periods of low external demands [21]. Likewise, functional networks associated with social, affective, mnemonic and executive systems are linked to different personality traits [41], which have been shown to relate to different patterns of ongoing thought [42, 43]. Furthermore, alterations in whole-brain connectivity have also been linked to a range of psychological disorders, like ADHD [44, 45], generalised anxiety [46], obsessive compulsive disorder [47, 48] and major depression ([49], see [50] for a review).

### Current study

This study set out to understand the relationship between trait variance in patterns of ongoing thoughts, dispositional features and neural organisation. We acquired retrospective descriptions of ongoing thoughts at rest after participants underwent a resting state functional magnetic resonance scan (rs-fMRI). In a separate session, we acquired measures of dispositional traits spanning both physical and mental health. We then calculated the functional connectivity of whole-brain functional networks and investigated how they related to ongoing cognition and psychological traits, as well as whether there are neural patterns common to both.

## Methods

### Participants

184 healthy participants were recruited by advert from the University of York. Written consent was obtained for all participants and the study was approved by the York Neuroimaging Centre Ethics Committee. 15 participants were excluded from analyses due to technical issues during the neuroimaging data acquisition or excessive movement during the fMRI scan (mean framewise displacement > 0.3 mm and/or more than 15% of their data affected by motion) [51], resulting in a final cohort of N = 169 (111 females, *μ_age_* = 20.1 years, *σ_age_* = 2.3).

### Behavioural methods

We asked participants to retrospectively report the experiences they had during the resting state fMRI scan, using a series of self-report questions. These items were measured using a 4-scale Likert scale, with the question order being randomised (all 25 questions are shown in Table 1). In order to identify the measures of experience with the best reliability and thus the most trait-like features, we repeated the resting state scanning session for a subset of our sample (N = 40) approximately 6 months later.

**Table 1.** Experience sampling questions asked at the end of the resting state fMRI scan.

| Dimension | Question<br>(My thoughts) |
| --- | --- |
| Vivid | ... were vivid as if I was there |
| Normal | ... were similar to thoughts I often have |
| Future | ... involved future events |
| Negative | ... were about something negative |
| Detail | ... were detailed and specific |
| Words | ... were in the form of words |
| Evolving | ... tended to evolve in a series of steps |
| Spontaneous | ... were spontaneous |
| Positive | ... were about something positive |
| Images | ... were in the form of images |
| Other | ... involved other people |
| Past | ... involved past events |
| Deliberate | ... were deliberate |
| Self | ... involved myself |
| Stop | ... were hard for me to stop |
| Distant time | ... were related to a more distant time |
| Abstract | ... were about ideas rather than events or objects |
| Decoupling | ... dragged my attention away from the external world |
| Important | ... were on topics that I care about |
| Intrusive | ... were intrusive |
| Problem solving | ... were about solutions to problems (or goals) |
| Here and now | ... were related to the here and now |
| Creative | ... gave me a new insight into something I have thought about before |
| Realistic | ... were about an event that has happened or could take place |
| Theme | ... at different points in time were all on the same theme |

To assess the participants’ physical and mental health, we administered well-established surveys at a later, separate session outside of the scanner. Quality of life, physical and psychological health, social relationships and environmental well-being were measured by the World Health Organization Quality of Life WHOQOL-BREF instrument [52]. Private and public self-consciousness and social anxiety were assessed using the Self-Consciousness scale [53], state and trait anxiety by the State-Trait Anxiety inventory [54] and trait rumination by the Ruminative response scale [55]. Finally, symptoms related to depression, autism, and ADHD were measured using the CES-D scale [56], the Autism Spectrum Quotient [57], and the World Health Organization Adult ADHD Self-Report scale [58] respectively. To reduce the dimensional structure of the measures of physical and mental health, we performed a principal component analysis (PCA) decomposition with varimax rotation.

### Neuroimaging methods

#### MRI data acquisition

MRI data were acquired on a GE 3 Tesla Signa Excite HDxMRI scanner, equipped with an eight-channel phased array head coil at York Neuroimaging Centre, University of York. For each participant, we acquired a sagittal isotropic 3D fast spoiled gradient-recalled echo T1-weighted structural scan (TR = 7.8 ms, TE = minimum full, flip angle = 20°, matrix = 256×256, voxel size = 1.13×1.13×1 mm^3^, FOV = 289×289 mm^2^). Resting-state functional MRI data based on blood oxygen level-dependent contrast images with fat saturation were acquired using a gradient single-shot echo-planar imaging sequence with the following parameters: TE = minimum full (≈19 ms), flip angle = 90°, matrix = 64×64, FOV = 192×192 mm^2^, voxel size = 3×3×3 mm^3^ TR = 3000 ms, 60 axial slices with no gap and slice thickness of 3 mm. Scan duration was 9 minutes which allowed us to collect 180 whole-brain volumes per participant.

#### fMRI data pre-processing

Functional MRI data pre-processing was performed using the Configurable Pipeline for the Analysis of Connectomes [59]. Pre-processing steps included motion correction by volume realignment (Friston 24-Parameter Model), nuisance signal regression of 24 motion parameters calculated in the previous step, plus five nuisance signals obtained by running a principal component analysis on white matter and cerebrospinal fluid signals using the CompCor approach [60], slice time correction, temporal filtering 0.009-0.1 Hz, spatial smoothing using a 6mm Full Width at Half Maximum of the Gaussian kernel and normalisation to MNI152 stereotactic space (2 mm isotropic) using linear and non-linear registration. No global signal regression was performed.

#### Group-ICA spatial maps

The resting state fMRI data were masked by a 20% probabilistic whole-brain grey matter mask, temporally demeaned, had variance normalisation applied and were fed into FSL’s MELODIC Incremental Group-PCA algorithm [61], and finally into group-ICA [62], where spatial-ICA was applied, resulting in 16 distinct group-ICA spatial maps. One component was marked as artefactual and removed. These group spatial maps were mapped onto each subject’s pre-processed data by running the first stage of a dual-regression analysis, which produced one time series per map per participant. The 15 spatial maps were named based on the top-loading term acquired from decoding each map on Neurosynth [63], and are shown in supplementary materials (Figure S1).

#### Static functional connectivity

Network modelling was carried out by using the FSLNets toolbox. We calculated the partial temporal correlation between the 15 ICA components’ time series, creating a 15 x 15 matrix of connectivity estimates for each participant and applied a small amount of L2 regularisation [64]. The connectivity values were converted from Pearson correlation scores into z-statistics with Fisher’s transformation (including an empirical correction for temporal autocorrelation). For group-level analyses, we combined all participants’ network matrices and ran a univariate general linear model, combined with permutation testing (FSL’s randomise, 5000 permutations) for each edge. Edge weight was the dependent variable and behavioural measures (thought questions or component loadings from the general and mental health PCA decomposition) were the independent variables. We used FWE-correction to account for multiple comparisons and all analyses controlled for age, gender and motion (mean frame-wise displacement) during the rs-fMRI scan.

## Results

### Describing experience and well-being

A test-retest reliability analysis revealed 6 questions, the responses to which were consistent within individuals across sessions (Fig. 1a): normal thoughts (i.e. experiences that I often have, intraclass correlation coefficient = 0.28, p = 0.035), deliberate vs spontaneous thoughts (ICC = 0.27, p = 0.044), intrusive thoughts (ICC = 0.29, p = 0.034), thoughts that took the form of attempts at problem-solving (ICC = 0.454, p = 0.001), thoughts about the here and now (ICC = 0.281, p = 0.036) and thoughts in the form of images (ICC = 0.3, p = 0.026). We used these items in subsequent analyses.

**Figure 1.**
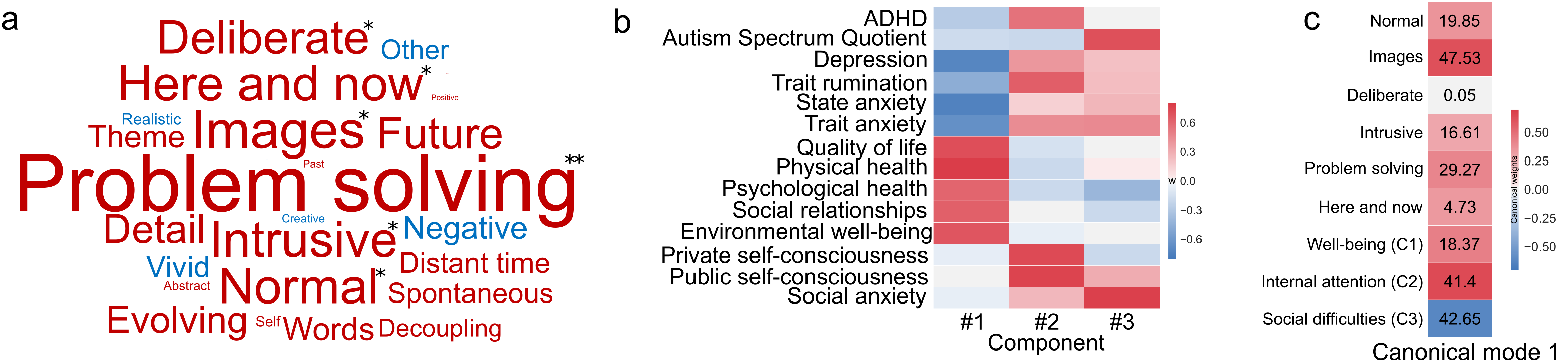
Behavioural variables and their relationship. **a**) Test-retest reliability of responses to items at the end of two resting state fMRI scans for 40 participants. Intraclass correlation coefficient (ICC) visualised using a word cloud. Font size represents the absolute ICC value and font colour its sign (red for positive and blue for negative values). * p < 0.05, ** p = 0.001. **b**) Component weights from a principal component analysis on the participants’ responses to dispositional measures of physical and mental health. **c**) Heat map showing the standardised canonical weights of each item for the significant canonical mode 1. The annotations indicate the item variance explained by the CCA mode.

The principal component analysis (PCA) of the measures of physical and mental health identified three principal components with eigenvalues greater than 1 (Fig. 1b). Component 1, *well-being*, loaded positively on measures of quality of life and psychological well-being and negatively on indices of depression and anxiety. Component 2, *internally directed attention*, loaded on self-consciousness, attention deficit hyperactivity disorder (ADHD) traits and rumination. Component 3, *social difficulties*, loaded on social anxiety and autism.

### Comparing common variance in disposition and descriptions of ongoing experience at rest

We used canonical correlation analysis (CCA) to describe the relationship between the PCA components of the physical and mental health scores and the six reliable self-report items. This yielded one significant canonical component, or mode, (F(18, 453.03) = 1.795, p = 0.023), explaining 11.5% of the shared variance (see Fig. 1c). The strongest experiential predictor was images (standardised canonical weight = 0.62, 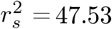) and the strongest trait predictors were patterns of internally directed attention (component 2, standardised canonical weight = 0.64, 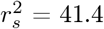) and a relative absence of social difficulties (component 3, standardised canonical weight = −0.62, 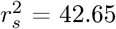).

### Comparing common variance in patterns of neural organisation and descriptions of ongoing experience

To understand the links between whole-brain connectivity and both ongoing experience and traits, we regressed the edges’ partial correlations against participants’ thoughts, while controlling for age, gender and motion during the rs-fMRI scan. We used multiple edge weight proportional thresholds (15%, 30%, 50% & no threshold) [65]. Taking into account only the strongest edges on average across subjects (top 15% of partial correlation weights, Fig. S2), we found more positive correlations between the precuneus and the lateral fronto-temporal network for subjects reporting thinking more in images (Fig. 2a) (p = 0.016, FWE-corrected). The pattern remained significant for a lower correlation weight threshold that included weaker edges in the analysis (top 30% of partial correlation weights) (p = 0.037, FWE-corrected). No significant relationship was found for a threshold that kept the top 50% of connections and when no threshold was applied at all.

**Figure 2.**
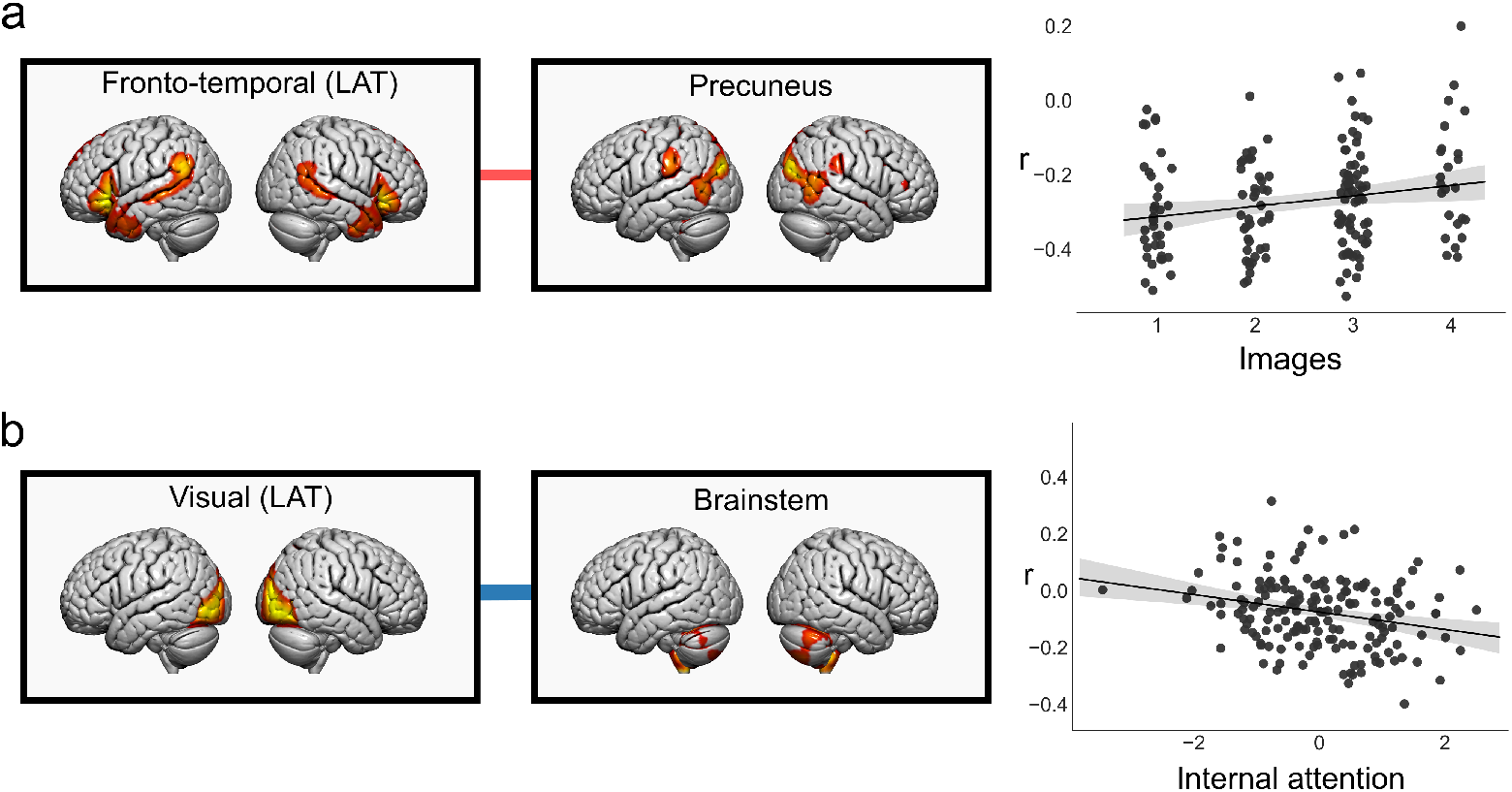
Static functional connectivity associated with behaviour. **a**) Left. The lateral frontotemporal and precuneus networks. The edge highlights an increase in their partial correlation for people thinking more in images. Right. Scatter plot of thinking in images scores with correlation values between these two networks for each participant. **b**) Left. The lateral visual and brainstem networks. The edge highlights a decrease in their partial correlation for higher internal attention scores (PCA Component 2). Right. Scatter plot of Component 2 scores from the PCA of the physical and mental health questionnaires with correlation values between these two networks for each participant.

### Comparing common variance in patterns of neural organisation and dispositional traits

Using the same connection thresholds, we found that a lower correlation between the lateral visual and the brainstem networks was associated with increased internally directed attention (PCA component 2) (Fig. 2b). The link was significant for the top 50% of connections (p = 0.021, FWE-corrected) and when no threshold was applied (p = 0.05, FWE-corrected).

### Examining the relationship between neural “fingerprints” of disposition and patterns of ongoing experience

Together, our analyses suggest that reports of imagery at rest were associated with traits of internally directed attention and that both features had reliable and distinct neural correlates. To understand if the identified neural patterns moderated the relationship between the psychological measures, we ran two multiple regression analyses; in one, the dependent variable was reports of visual imagery and in the other, dispositional internal focus (component 2, see Fig. 1). In each model, the remaining three scores (connectivity between the lateral fronto-temporal and precuneus networks, connectivity between the lateral visual and brainstem networks, and visual imagery or internal focus scores respectively) were entered as predictors, and we modelled their main effects and their three pairwise interactions (while controlling for age, gender, and motion during the resting state fMRI scan). The analysis with visual imagery as the dependent variable identified two main effects: thinking in images was positively correlated with the connectivity between the lateral fronto-temporal and precuneus networks (*β* = 0.19, p = 0.016) and with component 2 scores (*β* = 0.18, p = 0.026). Critically, it also indicated an interaction (*β* = 0.16, p = 0.043) suggesting that the correlations between the precuneus and lateral fronto-temporal regions and reports of imagery were stronger for participants who also had higher lateral visual and brainstem connectivity (associated with low loadings on internally directed attention). This is displayed in Figure 3 in the form of a scatter plot. The second analysis, with dispositional internal focus as the dependent variable, identified two main effects, but not an interaction: internally directed attention was negatively correlated with the connectivity between the lateral visual and brainstem networks (*β* = −0.25, p = 0.001) and positively correlated with thinking in images (*β* = 0.17, p = 0.024).

**Figure 3.**
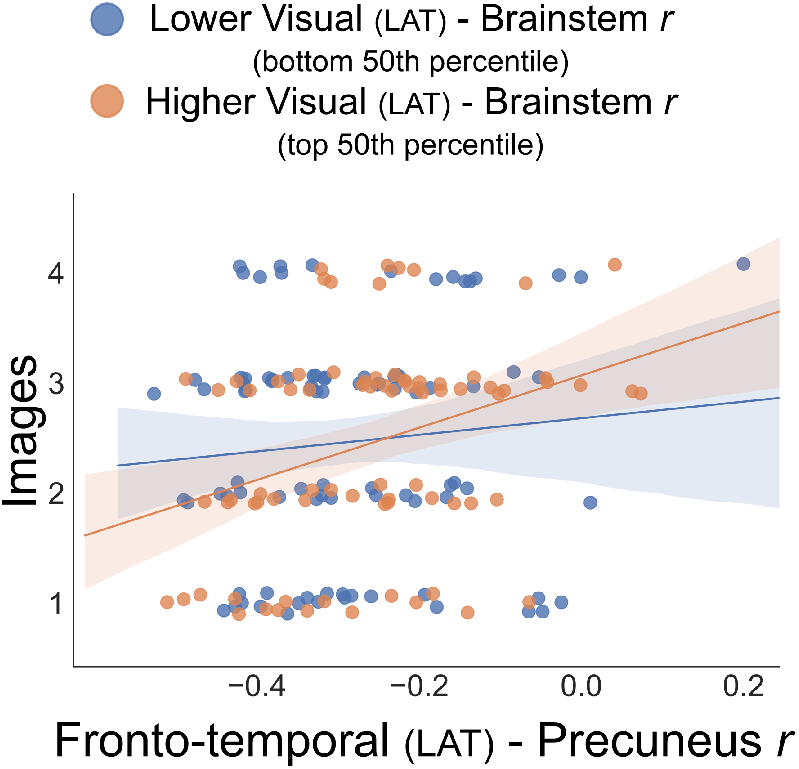
Moderation effect of the lateral visual - brainstem coupling on the relationship between thinking in images and lateral fronto-temporal - precuneus functional connectivity. The scatter plot shows the correlation values between the lateral fronto-temporal and precuneus networks and thinking in images scores grouped, by a median split, into higher and lower connectivity groups between the lateral visual and brainstem networks.

## Discussion

Our study set out to explore how associations between patterns of unconstrained ongoing experience and dispositional traits are reflected in the underlying neural architecture. We found a positive correlation between a style of thinking which emphasises visual imagery and a pattern of dispositional internal focus. Both of these psychological features had distinct neural correlates. Thinking more in images was related to an increase in correlation between the precuneus and a lateral fronto-temporal network, while a decreased correlation between the lateral visual and a brainstem/cerebellar network was linked to internal attention and ADHD traits.

Critically, we found a significant interaction between these two neural traits: the correlation between precuneus - lateral fronto-temporal connectivity and visual imagery was more pronounced for individuals who also had stronger coupling between the brainstem/cerebellar network and visual cortex. Our results, therefore, indicate that there are stable neural interactions that support ongoing experience as well as dispositional traits, and highlight that there are indirect neural relationships linking these phenomena together.

### Precuneus interactions with lateral frontal and temporal systems are important for visual imagery

Prior studies have shown that thinking in the form of images is an important feature of ongoing experience [66, 67] and our study suggests that this process is linked to a functional organisation in which a network anchored in the precuneus was more correlated with a network emphasising lateral regions of the temporal and frontal cortices. Although posterior medial cortex is functionally heterogeneous [68], the precuneus has been related to visuospatial mental imagery (for a review see [69]), episodic memory retrieval of imagined pictures [70] and imagining past episodes related to self [71]. Our analysis suggests that visual imagery is linked to functional connectivity between the precuneus and a lateral fronto-temporal network that contributes to multi-modal forms of semantic cognition [72]. Contemporary accounts suggest that the anterior temporal lobe supports cognition with different representational features (pictures versus words, [73] for a review). Generally therefore, our results are consistent with the hypothesis that the production of visual imagery depends upon the integration of a system important for visual processing (the precuneus) with regions supporting multi-modal elements of cognition (lateral temporal and frontal cortex).

### Lateral visual cortex and brainstem interactions relate to rumination, self-consciousness and ADHD traits

Rumination, self-consciousness and ADHD characteristics loaded on the same component derived from the PCA decomposition of the questionnaire data, in line with previous studies investigating their relationships directly [74, 75] or through their shared characteristics with different and often maladaptive types of self-generated thought [22, 24, 76, 77]. Our neuroimaging analyses showed that this behavioural component was linked to reduced correlation between the brainstem/cerebellum and the lateral visual cortex. Previous studies have shown that brainstem abnormalities can support increased accuracy of diagnostic classification in ADHD [78]. Decreased axial diffusivity (a marker of axonal degeneration in clinical cases, [79]) in the occipital lobe and brainstem has also been associated with an ADHD diagnosis [80]. In addition, processes within the brainstem have been related to rumination, worry and anxiety by influencing heart rate variability [81] and by its functional interactions with the limbic network in anxiety disorders [82]. Links between increased rumination, self-consciousness and ADHD traits may highlight a functional neural configuration where visual input becomes increasingly incoherent with the activity of a network regulating autonomic functions and modulating global connectivity [83], potentially reflecting the reduced processing of sensory input that is important for internal focused experience [84].

### Interactions between neural correlates of disposition and patterns of ongoing thought

Given the results from our behavioural analyses (see Fig. 1) and multiple evidence linking ongoing thought and dispositional traits, we also investigated whether there was an interaction between their unique neural correlates. Notably, our analyses highlighted that the association between precuneus-lateral fronto-temporal connectivity and reports of visual imagery was more pronounced for individuals who had stronger connectivity between regions of the brainstem and lateral occipital cortex. We suggest this could reflect a more integrated global brain state, given that contemporary views of the basal ganglia, cerebellum, and the cortex emphasise that these discrete systems should be seen as parallel features of an integrated system [85]. We speculate that stronger lateral visual, brainstem coupling gives rise to more stable neural conditions, resulting in an increase in integration between precuneal and lateral frontal and temporal systems, and thus facilitating the production of visual imagery. More generally, our study suggests that understanding the neural mechanisms underlying patterns of dispositional traits may be improved by accounting for features of ongoing experiences that emerge when neural activity is recorded (and vice versa). As well as the questionnaire administered in our study, there is now an emerging number of measures of ongoing experience such as the New York Cognition Questionnaire [5], the Amsterdam Resting State Questionnaire [86] and Resting state questionnaire [67]. The low economic cost of these measures and the ease of their administration suggest that collecting data on patterns of ongoing thought can improve our understanding of neural correlates of specific dispositional states. We recommend that these measures become part of standard resting state protocols.

### Limitations

Although our study highlights the complex relationship between dispositional traits, patterns of ongoing thought and measures of functional connectivity, it leaves several important questions unanswered. First and foremost, it is unclear why our analyses identified correlates for imagery rather than the other self-report items we studied. As well as issues with experimental power, it is likely that this was partly because we lack an agreed upon ontology of ongoing thought upon which the selection of the appropriate self-report items can be made. This means that there may well be aspects of ongoing cognition that we failed to measure, and others that the wording of our questions may have made the detection of neural correlates more difficult. Thus, although our study suggests that there are likely to be important relationships between ongoing thought patterns, dispositional traits and functional connectivity, it remains an important open question what categories of experience are most important to study and with which self-report measures they should be assessed. In addition, we measured ongoing thoughts retrospectively, an approach that is suitable for trait level inferences, but less useful for detecting neural correlates which are short and/or occur infrequently [84, 87]. It is possible that our measure of brain activity was not optimal for determining associations with certain types of experience. It may be that using metrics sensitive to dynamic changes in neural activity [88] would increase our capacity to identify neural correlates other than those identified in this study. Finally, it is important to note that patterns of ongoing experience are at least partly context dependent [42, 89]. Although the resting state is a well-recognised technique in the cognitive neuroscientists toolbox, it is possible that the neural correlates of certain aspects of experience may be detected more readily in a different tasks environment. For example, the neural correlates of both patterns of off-task thought and detailed task focus were easier to determine using machine learning during a demanding 1-back task [4]. It is also important to note that our sample was of young university undergraduate and post graduate students and measures of psychopathology and other disorders within such a population may not generalise fully to clinical groups. Extending this work to clinical conditions, especially those which are partly identifiable through their patterns of cognition, will be important. For example, studies have found that patterns of repetitive verbal thoughts can be a characteristic of anxiety [90]. Moving forward, it will be important to develop better self-report measures, more generalisable populations and to use alternative techniques for mapping neural activity to completely understand the neural correlates of different aspects of ongoing thought.

### Conclusion

In conclusion, our study highlights that neural functioning at rest provides information related to ongoing experience, patterns of dispositional traits, and their association. Our test-retest analyses revealed recurring themes of spontaneous thoughts that remain relatively stable within individuals. We found that thoughts based on imagery were related to stronger interactions between a network anchored in the precuneus and lateral temporal regions of the temporal and frontal cortex, many of which are members of the DMN. More generally, our findings complement previous studies investigating the neural substrates of ongoing experience and suggest that the impact of the functional interactions of the brain on the human condition can be better understood by accounting for both dispositional traits and patterns of ongoing experience.

## Data availability

The data that support the findings of this study are available from the corresponding author upon reasonable request.

## Acknowledgments

This work was supported by European Research Council awarded to JS (WANDERINGMINDS - 646927). The authors would like to thank Giulia Poerio, Deniz Vatansever, Mladen Sormaz, Charlotte Murphy and Hao-Ting Wang for their help.

