## Supplementary Material for "Interactions between the neural correlates of dispositional internally directed thought and visual imagery"

### Supplementary Information

#### ICA decomposition of the fMRI data

We performed an independent component analysis (ICA) on the preprocessed concatenated fMRI data and opted for a 16-component solution. On visual inspection of the derived components, one was marked as artefactual, while the rest resembled well-known whole-brain functional networks. The 15 spatial maps were named based on the top-loading term acquired from decoding each map on Neurosynth (`https://neurosynth.org`) and are shown in Figure 1.

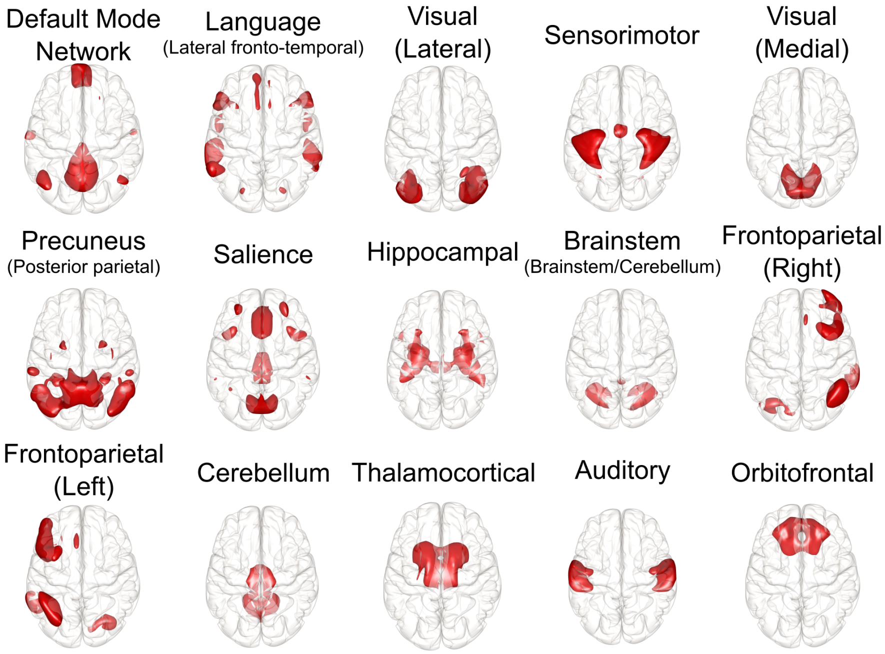


Figure 1: *ICA decomposition of fMRI data.* The panel shows the 15 spatial maps obtained from an independent component analysis on the temporally concatenated fMRI data. The maps correspond to well-known whole-brain functional networks and were named based on the top-loading term acquired from decoding each map on Neurosynth.

#### Functional connectivity of ICA components

Figure 2 shows the average partial correlation matrix across subjects (thresholded at the top 15% of partial correlation weights) in the form of a chord plot. Strong interactions highlight the relatively decreased correlation between the DMN and sensory networks, the auditory with the sensorimotor and the two visual networks forming synchronised pairs of sensory processing, and the precuneus with the salience networks as positively coupled.

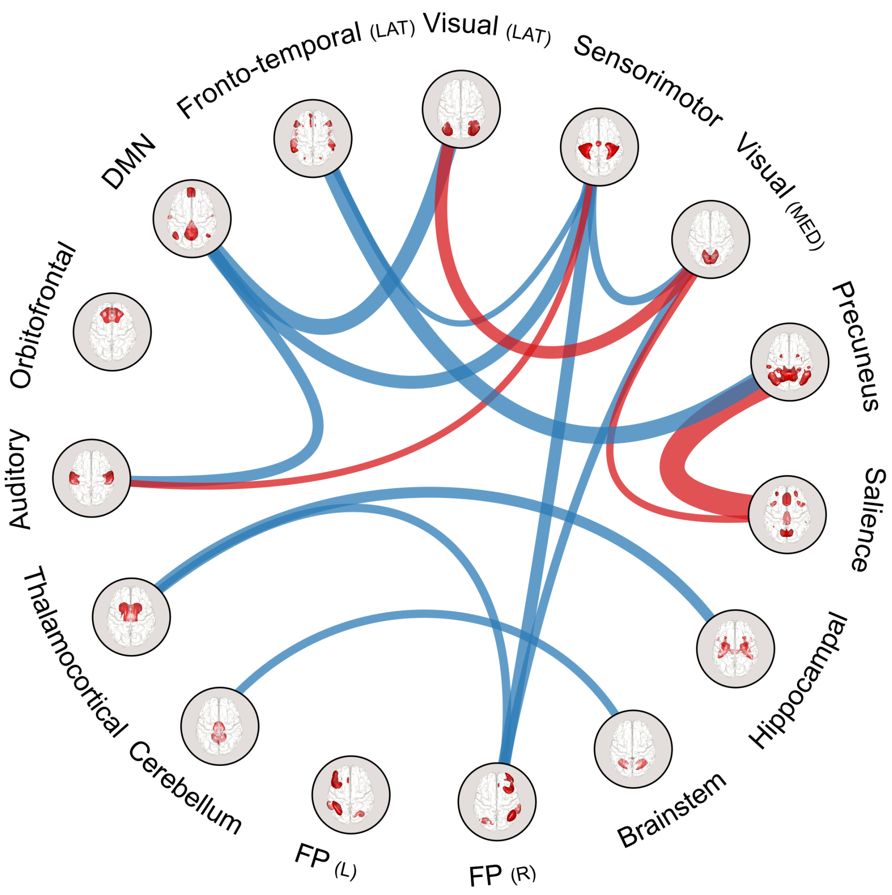


Figure 2: *Partial correlation of ICA maps.* Network circle graph of the average partial correlation matrix across subjects. The circle graph was thresholded at the $85^{th}$ percentile of absolute z-values as estimated from an r to z Fisher's transformation. Red is for positive and blue for negative values and edge width is weighted by their absolute value. L:Left, R:Right, LAT:Lateral, MED:Medial.
